# Multivariate Neural Patterns of Reward and Anxiety in Adolescents with Anorexia Nervosa

**DOI:** 10.1101/2024.07.23.604826

**Authors:** Hayden J. Peel, Nicco Reggente, Michael Strober, Jamie D. Feusner

## Abstract

People with anorexia nervosa (AN) commonly exhibit elevated anxiety and atypical reward responsiveness. To examine multivariate neural patterns associated with reward and the impact of anxiety on reward, we analyzed fMRI data from a monetary reward task using representational similarity analysis, a multivariate approach that measures trial-by-trial consistency of neural responses. Twenty-five adolescent girls with AN and 22 mildly anxious controls lacking any history of AN were presented personalized anxiety-provoking or neutral words before receiving a reward, and neural response patterns in reward regions were analyzed. Consistent with our preregistered hypothesis, AN participants showed lower representational similarity than controls during neutral-word rewarded trials. Within groups, controls showed significant representational similarity in reward circuit regions including the left nucleus accumbens, left basolateral amygdala, and left medial orbitofrontal cortex, which were not observed in AN. Further, reward-related prefrontal cognitive control areas – left ventrolateral prefrontal cortex and left dorsolateral prefrontal cortex – showed significant representational similarity in both groups, but a larger spatial extent in controls. Contrary to predictions, there were no significant between-group differences for the effects of anxiety-words on reward representational similarity, and representational similarity did not predict longitudinal symptom change over six months. Overall, the results demonstrate relatively inconsistent trial-by-trial responses to reward receipt in the neutral state in AN compared with controls in both reward circuit and cognitive control regions, but no significant differential effects of anxiety states on reward responses. These results add to dynamic understandings of reward processing in AN that have potential implications for planning and guiding reward-focused interventions.

## 1 Introduction

Anorexia nervosa (AN) is an eating disorder characterized by restrictive food intake, fear of gaining weight, and a disturbed body image[1], with one of the highest mortality rates in psychiatry[2, 3]. Anxiety is near-universally present before weight loss and during early disordered eating [4–6], contributing to behaviors like restrictive eating, food avoidance, excessive exercise, and body checking. Some behaviors, such as meeting a weight goal, or being able to feel the protrusion of one’s hip bones, may be experienced as rewarding. Conversely, typically rewarding experiences—such as consuming palatable foods or experiencing prosocial emotions—are less subjectively pleasant in AN compared to controls (reviewed in [7]). Thus, both negative reinforcement of avoiding anxiety-inducing stimuli and positive reinforcement of rewarding feelings generated by meeting one’s food, weight, or body goals may be operating. Yet, little is known of how neural systems involved in anxiety and reward directly interact in those with AN, which may have clinical relevance in the onset or maintenance of AN symptoms.

Mass-univariate neuroimaging studies in AN have revealed distinct brain activity patterns in key reward and anxiety circuits compared to healthy controls and those with other eating disorders. Reward processing involves anticipation, experiencing, and learning [8], and dysregulation in these domains have been reported in AN [9]. Reward receipt (the focus of this study), typically involves the ventral tegmental area (VTA), striatum, and orbitofrontal cortex [8]. Individuals with AN show different activity patterns in these areas, including lower activation in the VTA and dorsal striatum in response to palatable food cues compared to those with bulimia nervosa [10]. Although not a traditional rewarding stimulus to those without AN, women with AN exhibit higher activation in the ventral striatal reward system when processing underweight stimuli compared to normal-weight stimuli, whereas healthy controls show the opposite pattern [11]. AN also affects social reward processing, with reduced activation in regions like the medial prefrontal cortex, striatum, and nucleus accumbens (NAcc) [12]. Differences in anxiety circuitry are also well documented, with increased activity in the medial orbitofrontal cortex (mOFC), insula, anterior cingulate cortex, amygdala-hippocampal area, and medial/lateral prefrontal regions, and decreased activity in the cerebellum in response to anxiety-inducing stimuli related to food or body image [13–17].

These univariate studies demonstrate different degrees of activity in brain areas but may miss subtler, distributed patterns of neural activity that underlie complex psychological states. A multivariate approach, like representational similarity analysis (RSA) [18] examines activity across multiple voxels or brain areas, and quantifies pattern similarity or dissimilarity. By capturing distributed neural representations of anxiety and reward, RSA may reveal how different aspects of these states interact and influence each other in terms of response consistency. For instance, RSA could identify how the neural representation of reward might change when a person is anxious versus not, focusing on patterns and representational space changes beyond activation degree. High representational similarity indicates a consistent, coordinated multivariate response, likely reflecting longstanding patterns of repeated activation or conditioned responses [19]. Given the self-reinforcing nature of anxiety and reward symptoms in AN, understanding the consistency of brain response patterns with RSA is crucial; yet, our knowledge of multivariate patterns in anxiety and reward circuitry in AN is limited.

Our previous work [20] found higher representational similarity in AN for anxiety-provoking compared to neutral stimuli in prefrontal regions, including the frontal pole, medial prefrontal cortex, dorsolateral prefrontal cortex (dlPFC), and mOFC. These results clarify which nodes within a larger anxiety network are contributing to consistent responses to anxiety-inducing word stimuli, supporting theories that altered mechanisms of cognitive control could be engaged by frontal brain areas in AN in an anxious state [21]. To the authors’ knowledge, no experiments have investigated multivariate reward patterns in AN or how these may be influenced differentially from healthy controls in anxious or neutral states.

The present study aimed to investigate multivariate reward and anxiety patterns in AN with RSA. Hypotheses were pre-registered [https://aspredicted.org/F3D_418]. Participants completed a task in which they either received a reward (reward trials) or did not (non-reward trials), with the reward preceded by either a personalized anxiety-evoking or neutral word (see Methods for further details). Our first hypothesis was that in those with AN, reward response following neutral states would be more variable, and thus less consistent (i.e., lower representational similarity; RS) in reward regions. We reasoned that previous studies showing differences between AN and controls as described above in reward responses might be accounted for by reduced consistency (greater heterogeneity) in brain patterns. Second, we investigated whether anxiety-induced states affect between-group differences in reward receipt. Previous research has suggested that high anxiety might ‘hijack’ the reward response, a concept explored in various contexts [22–24], but which has not been tested in AN. Given the high salience of disorder-specific anxiety-inducing stimuli in AN [9], and previously found consistent responses to anxiety provocation [20], we hypothesized that AN individuals would exhibit more consistent reward responses. This would be due to consistent effects trial-by-trial of induced anxiety on reward responses, reflected in higher RS in reward regions in AN compared to controls. We also considered whether these multivariate patterns demonstrate clinical relationships. We hypothesized that neutral state RS would be associated with poorer behavioral activation system (BAS) reward responsiveness. We also hypothesized that the difference in RS between anxiety and neutral conditions in reward regions following anxiety provocation (i.e., the additional effect of anxiety on reward responses) would predict worse clinical course over six months after intensive treatment, indicated by a decrease in body mass index (BMI) and an increase in eating disorder symptom severity scores.

## 2 Materials and Methods

### 2.1 Participants

The UCLA Institutional Review Board approved the study. Females aged 10-19 were recruited from the greater Los Angeles area. Written informed consent was obtained from all participants, and from parents or legal guardians for those under 18. Recruitment for controls was conducted through online and community advertisements and flyers. AN participants met DSM-5 criteria (other than needing to currently be at a significantly low body weight) and were recruited from the UCLA inpatient eating disorder unit and local treatment centers, enrolled at the end of their treatment once they met the hospital or treatment center criteria for transitioning to a lower level of care, and were either partially or fully weight-restored. Non-clinical but mildly anxious controls were included if they did not meet criteria for any DSM-5 disorder, did not take any medications, but scored at least 0.5 standard deviations above population norms on the anxiety portion of the DASS-21. (See Supplementary Materials S1-S2 for full inclusion and exclusion criteria, as well as clinical assessments.) The study was registered on Clinicaltrials.gov NCT02948452.

### 2.3 fMRI task paradigm

Participants completed a monetary reward paradigm in a fast event-related design. Briefly, they classified colored fractal images into two groups, with the potential to earn $10 for correct answers. After classification with a button press, a word was presented. The word presented differentiated the two tasks completed at two time points, two days apart in a randomized order. One version used personalized anxiety-provoking words, and the other used neutral words. Detailed word selection and event timing information are in the Supplementary Materials S6 and are described here [25]. Following the word presentation, feedback was given about receiving or not receiving a reward (50% were randomly rewarded), which was the main period of interest. There were 60 randomized trials in total, resulting in approximately 30 reward trials of interest per participant: anxiety-word rewarded trials, neutral-word rewarded trials, anxiety-word non-rewarded trials, and neutral-word non-rewarded trials.

### 2.4 MRI data acquisition and processing

MRI data were acquired on a 3T Siemens PRISMA scanner. Functional MRI data preprocessing was done with FSL (FMRIB’s Software Library, www.fmrib.ox.ac.uk/fsl) using FEAT (FMRI Expert Analysis Tool). See Supplementary Material S4-S5 for details of data acquisition and processing, including quality control and motion correction.

### 2.5 Regions of interest

Core areas involved in reward comprised the reward ROI mask, including bilateral basolateral amygdala, VTA, NAcc, mOFC, as well as regions involved in reward-related cognitive control - dorsolateral prefrontal cortex (dlPFC), left ventrolateral prefrontal cortex (vlPFC), and supplementary motor area (SMA) [26]. See Supplementary Materials S7 for details on how these areas were defined.

### 2.7 Representational similarity analysis

We conducted RSA within the ROIs using custom MATLAB scripts to create RSA maps for each participant for the anxiety-word rewarded trials, neutral-word rewarded trials, anxiety-word non-rewarded trials, and neutral-word non-rewarded trials. We also had pre-registered an exploratory analysis, testing the hypothesis that a reward state may influence a subsequent anxiety state. For more details on this analysis, see Supplementary Materials S8. These maps were generated by correlating single-trial beta maps from the reward and anxiety word trials. A searchlight mapping approach [27] within the ROI masks was also used, creating spherical ROIs with a 2-voxel radius centered on each voxel. We calculated the mean Pearson correlation across all trials within each condition for each spherical ROI. This resulted in similarity maps for each period for each participant, where each voxel’s value represented the average correlation (r-value) across all trials when the spherical ROI was centered on that voxel. Scripts used to conduct these analyses are available open-source (https://github.com/Institute-for-Advanced-Consciousness/Anorexia-Reward-RSA).

### 2.8 Statistical analysis

#### Group Analyses

##### Averaged ROI analysis

To compare multivariate responses across the reward mask, we r-to-z transformed and averaged the RS values, resulting in a single z-value per participant for each condition and ROI [28]. These values were analyzed using ANCOVAs with group as the factor, pubertal development scale (PDS) scores as a covariate, and *p* < .05, two-tailed, as the statistical threshold. As a planned analysis to determine if any specific region within the mask contributed more significantly to the results, we performed MANOVA with group as a factor, PDS as a covariate, and all 7 reward/reward-related cognitive control ROIs as the multivariate dependent variables; we then iteratively removed each region of the ROI and re-calculated the MANOVA to check for changes in the results.

##### Searchlight analysis

Within the ROI, group differences were assessed with FSL, including PDS values as a covariate of non-interest. We conducted a non-parametric permutation test with 5000 permutations and a threshold-free cluster enhancement method [29] using FSL’s Randomise, applying a statistical threshold of *p* < .05, FWE-corrected. As a post hoc analysis, within-group responses for various conditions were tested using one-sample t-tests with PDS as a covariate.

#### Associations with clinical variables

A series of linear mixed models (LMM) were performed using data from the AN group only. All used PDS as a covariate. The first model included adjusted-BMI (i.e., BMI-percentiles) as the dependent variable, with time (adjusted-BMI entry, and monthly intervals to 6-month follow-up) and ‘across anxiety and reward’ RS as factors. The ‘across anxiety and reward’ RS was computed by subtracting the beta maps of single-trial neural activity during neutral reward conditions from those during anxiety reward conditions. This metric was utilized in the LMMs to examine RS that specifically correlates with the reward response, independently of any RS differences linked to preceding anxiety or neutral word stimuli. The second LMM was similar but used eating disorder examination (EDE) global scores as the dependent variable and included only two time points (entry and 6-month follow-up). The final LMM used BAS as the dependent variable, with time (BAS entry and 6-month follow-up) and RS values from the neutral-word reward period as factors (but did not specify any interaction term).

Post hoc exploratory Pearson’s and Spearman’s partial correlations (with PDS as a covariate) were used to examine associations between symptom severity measures and RS that differed between groups in the ANCOVA. To reduce the chance of false discoveries, we applied the Benjamini-Hochberg False Discovery Rate (FDR) correction to the *p*-values from the correlation analyses.

## 3 Results

### 3.1 Sample characteristics

Twenty-five females with AN (14.6 ± 1.9 years) and 22 control participants (16.2 ± 1.8) were included in analyses. Mean age in months (*p* = .005), PDS (*p* < .001), and BMI (*p* <. 001) were lower in AN. The AN group had significantly higher Hamilton Anxiety Rating Scale (HAM-A), Depression Anxiety Stress Scale (DASS) depression, Children’s Depression Rating Scale (CDRS), Eating Disorder Examination (EDE) global, and Yale-Brown-Cornell Eating Disorder Scale (YBC-EDS) scores (all *p* < .001). No differences were found for DASS-anxiety (*p* = .977) or BAS scores (*p* = .669). Detailed information can be found in Table 1 below.

**Table 1.**
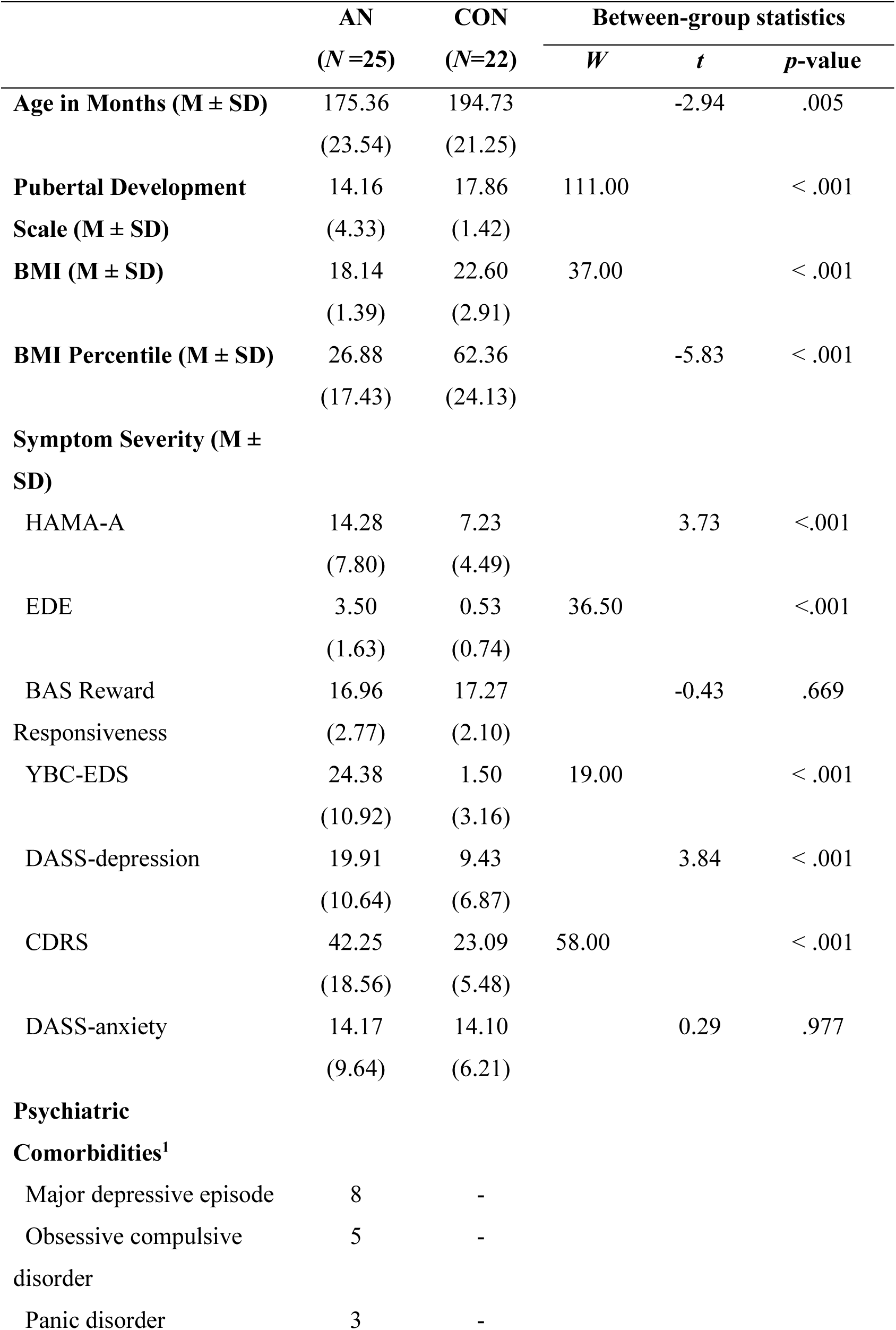
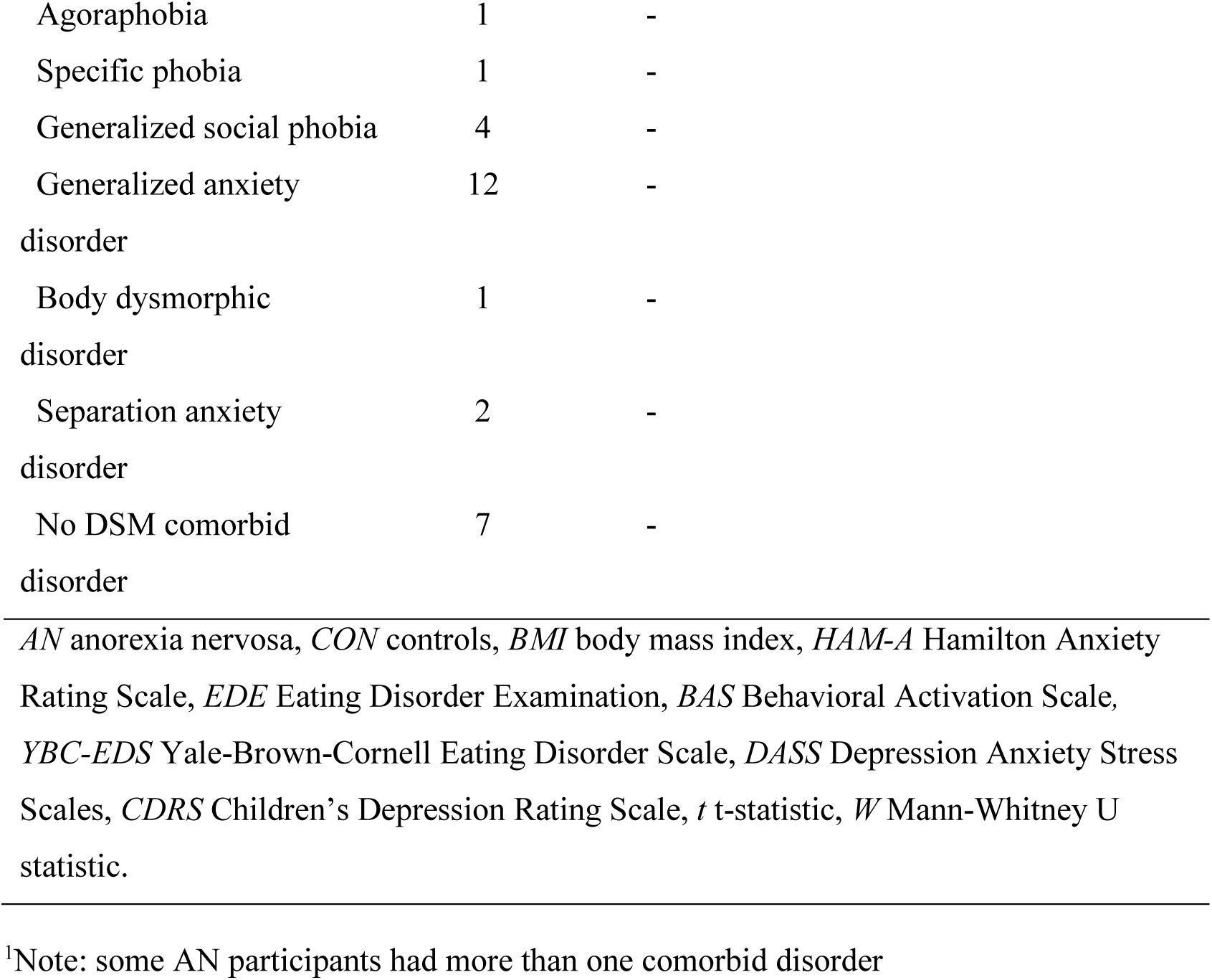
Demographics and Psychometrics.

|  | AN<br>(*N* =25) | CON<br>(*N*=22) | Between-group statistics |  |  |
| --- | --- | --- | --- | --- | --- |
|  |  |  | *W* | *t* | *p*-value |
| **Age in Months (M ± SD)** | 175.36<br>(23.54) | 194.73<br>(21.25) |  | -2.94 | .005 |
| **Pubertal Development Scale (M ± SD)** | 14.16<br>(4.33) | 17.86<br>(1.42) | 111.00 |  | < .001 |
| **BMI (M ± SD)** | 18.14<br>(1.39) | 22.60<br>(2.91) | 37.00 |  | < .001 |
| **BMI Percentile (M ± SD)** | 26.88<br>(17.43) | 62.36<br>(24.13) |  | -5.83 | < .001 |
| **Symptom Severity (M ± SD)** |  |  |  |  |  |
| HAMA-A | 14.28<br>(7.80) | 7.23<br>(4.49) |  | 3.73 | <.001 |
| EDE | 3.50<br>(1.63) | 0.53<br>(0.74) | 36.50 |  | <.001 |
| BAS Reward Responsiveness | 16.96<br>(2.77) | 17.27<br>(2.10) |  | -0.43 | .669 |
| YBC-EDS | 24.38<br>(10.92) | 1.50<br>(3.16) | 19.00 |  | < .001 |
| DASS-depression | 19.91<br>(10.64) | 9.43<br>(6.87) |  | 3.84 | < .001 |
| CDRS | 42.25<br>(18.56) | 23.09<br>(5.48) | 58.00 |  | < .001 |
| DASS-anxiety | 14.17<br>(9.64) | 14.10<br>(6.21) |  | 0.29 | .977 |
| **Psychiatric Comorbidities<sup>1</sup>** |  |  |  |  |  |
| Major depressive episode | 8 | - |  |  |  |
| Obsessive compulsive disorder | 5 | - |  |  |  |
| Panic disorder | 3 | - |  |  |  |

|  |  |  |
| --- | --- | --- |
| Agoraphobia | 1 | - |
| Specific phobia | 1 | - |
| Generalized social phobia | 4 | - |
| Generalized anxiety disorder | 12 | - |
| Body dysmorphic disorder | 1 | - |
| Separation anxiety disorder | 2 | - |
| No DSM comorbid disorder | 7 | - |

*AN* anorexia nervosa, *CON* controls, *BMI* body mass index, *HAM-A* Hamilton Anxiety Rating Scale, *EDE* Eating Disorder Examination, *BAS* Behavioral Activation Scale, *YBC-EDS* Yale-Brown-Cornell Eating Disorder Scale, *DASS* Depression Anxiety Stress Scales, *CDRS* Children's Depression Rating Scale, *t* t-statistic, *W* Mann-Whitney U statistic.
<sup>1</sup>Note: some AN participants had more than one comorbid disorder

### 3.2 Between- and within-group results

ANCOVA revealed a significant difference between groups for the neutral-word rewarded trials (*F* _(1, 44)_ = 6.44, *p* = .015), driven by lower RS in the AN group compared to controls in the ROI mask. Iteratively removing each region from the reward ROI mask and performing MANOVA did not find any significant differences for different combinations of ROIs (all *p* > .148), suggesting that the between-group results were not heavily influenced by any one region within the overall reward mask. Between-group searchlight analyses within the ROI mask did not reveal significant differences between groups, however, within-group results showed significant clusters for the control group which had much larger spatial extents or were not observed in the AN group. Subcortical reward circuit areas not observed in AN included the left NAcc and left basolateral amygdala, as well as left mOFC. Reward-related cognitive control areas that appeared in both groups and had a similar magnitude but a smaller spatial extent in AN included left vlPFC and left dlPFC, as well as right mOFC (see Figure 1 and Table 2).

**Figure 1.**
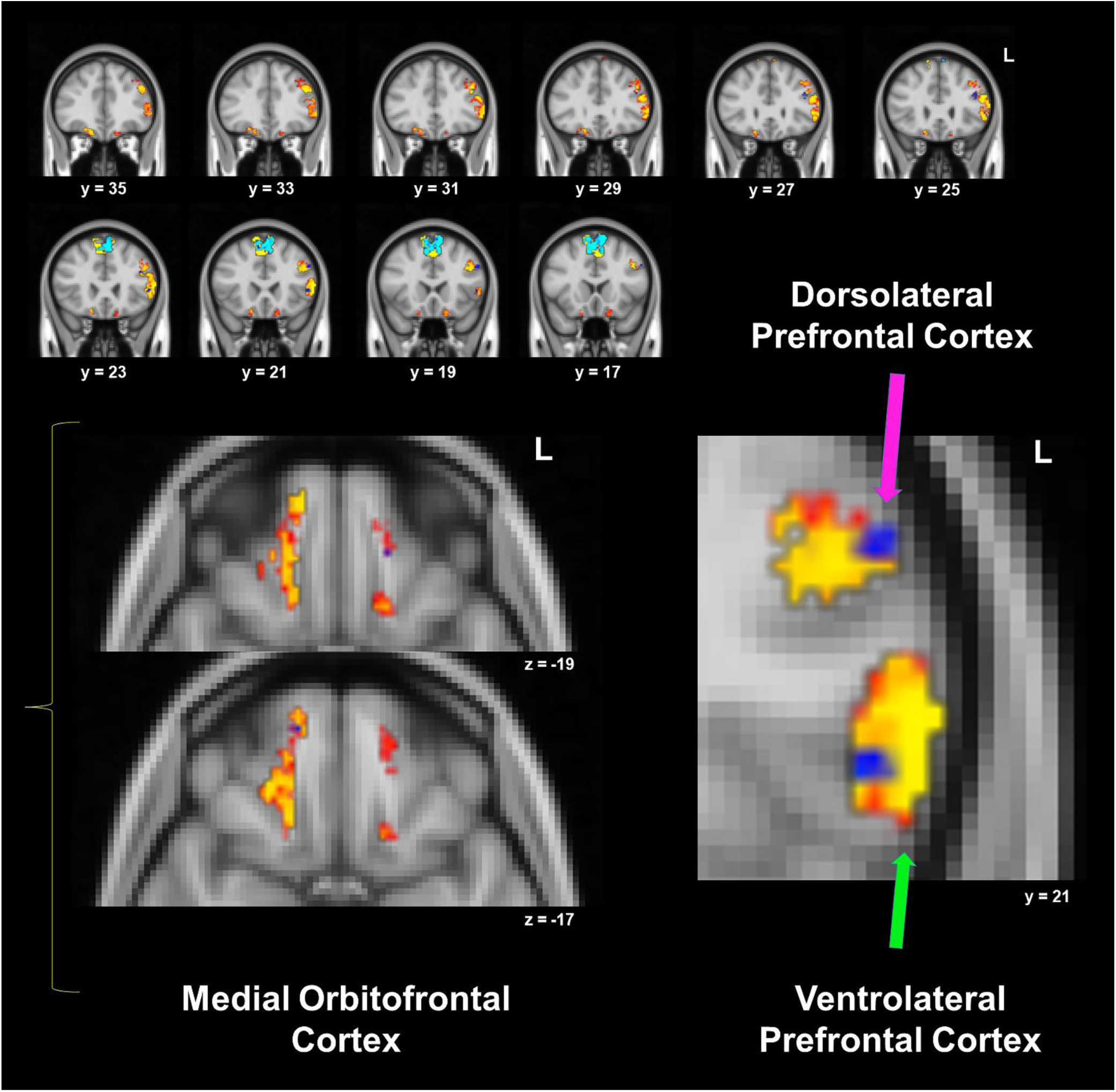
Within-groups one sample *t*-tests for neutral-word rewarded trials’ representational similarity, from the searchlight analysis within the mask that included reward and reward-related cognitive control regions. Control and anorexia nervosa (AN) within-group results are overlaid: red/yellow clusters depict significant areas in controls, while blue clusters are significant in AN (*p* < .05, corrected). See Table 2 for details for each cluster.

**Table 2.** Within-group one-sample *t* – test for neutral-word rewarded trials for AN and controls.

| Region | Size<br>(voxels) | T max | Cluster # | MNI Coordinates |  |  |
| --- | --- | --- | --- | --- | --- | --- |
|  |  |  |  | X | Y | Z |
| **Control Group** |  |  |  |  |  |  |
| **Supplementary Motor Area** | 2862 | 8.62 | 8 | -2 | 16 | 16 |
| **L vIPFC/dIPFC** | 806 | 6.58 | 7 | -54 | 22 | 8 |
| **R mOFC** | 232 | 6.75 | 6 | -14 | 54 | -20 |
| **L mOFC** | 65 | 5.97 | 5 | -10 | 38 | -24 |
| **L mOFC** | 44 | 6.90 | 4 | -16 | 20 | -20 |
| **L Basolateral Amygdala** | 2 | 6.95 | 3 | -28 | -2 | -28 |
| **L NAcc** | 2 | 8.66 | 2 | -6 | 8 | -8 |
| **AN Group** |  |  |  |  |  |  |
| **Supplementary Motor Area** | 2122 | 8.22 | 7 | 8 | -4 | 48 |
| **L dIPFC** | 33 | 6.42 | 6 | -40 | 28 | 24 |
| **L vIPFC** | 23 | 6.03 | 5 | -48 | 22 | 4 |
| **L dIPFC** | 19 | 5.61 | 4 | -52 | 20 | 32 |
| **R mOFC** | 4 | 6.61 | 3 | 14 | 54 | -16 |
| **L dIPFC** | 4 | 4.67 | 2 | -42 | 22 | 32 |

AN anorexia nervosa, *L* left hemisphere, *R* right hemisphere, *vIPFC* ventrolateral prefrontal cortex, *dIPFC* dorsolateral prefrontal cortex, *mOFC* medial orbitofrontal cortex, *NAcc* nucleus accumbens, *MNI* Montreal Neurological Institute

ANCOVA found no significant group differences in RS during anxiety-word rewarded trials (*F* _(1, 44)_ = 1.72, *p* = .197). No group differences were detected for control analyses: non-rewarded neutral-word trials (*F* _(1, 44)_ = 0.97, *p* = .333) and non-rewarded anxiety-word trials (*F* _(1, 44)_ = 0.39, *p* = .536). Similarly, the between-group searchlight analyses did not reveal any significant differences between groups for each trial type, and MANOVA did not find that any one region contributed more strongly to results. However, within-group searchlights yielded some notable outcomes. For the anxiety-word rewarded trials, controls had significant RS clusters in the left mOFC and left vlPFC that were not present in AN (Figure 2A, Table 3). For the neutral-word non-rewarded trials, right dlPFC and left basolateral amygdala were observed in controls but not AN. For the anxiety-word non-rewarded trials, significant right basolateral amygdala RS was detected in AN but not controls, while clusters in the left basolateral amygdala, right NAcc, and left mOFC were significant in controls but not AN. For the control ‘non-rewarded’ trial results, see Supplementary Materials Figure S1 and Tables S1 and S2.

**Figure 2.**
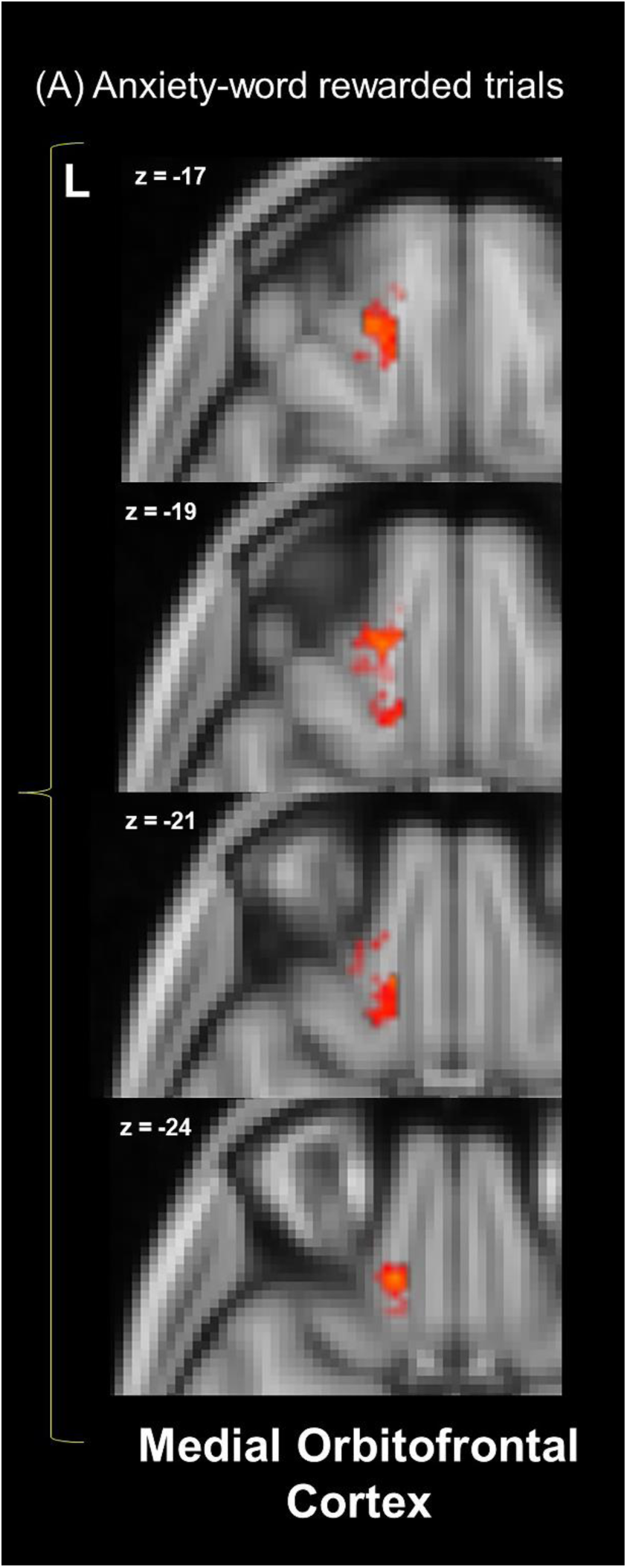
Within-group one-sample *t*-tests for the (A) anxiety-word rewarded trials representational similarity from the searchlight analysis within the reward mask. Within-group results are overlaid: red/yellow clusters depict significant areas in controls, while there are no significant blue clusters in AN (*p* < .05, corrected).

**Table 3.**
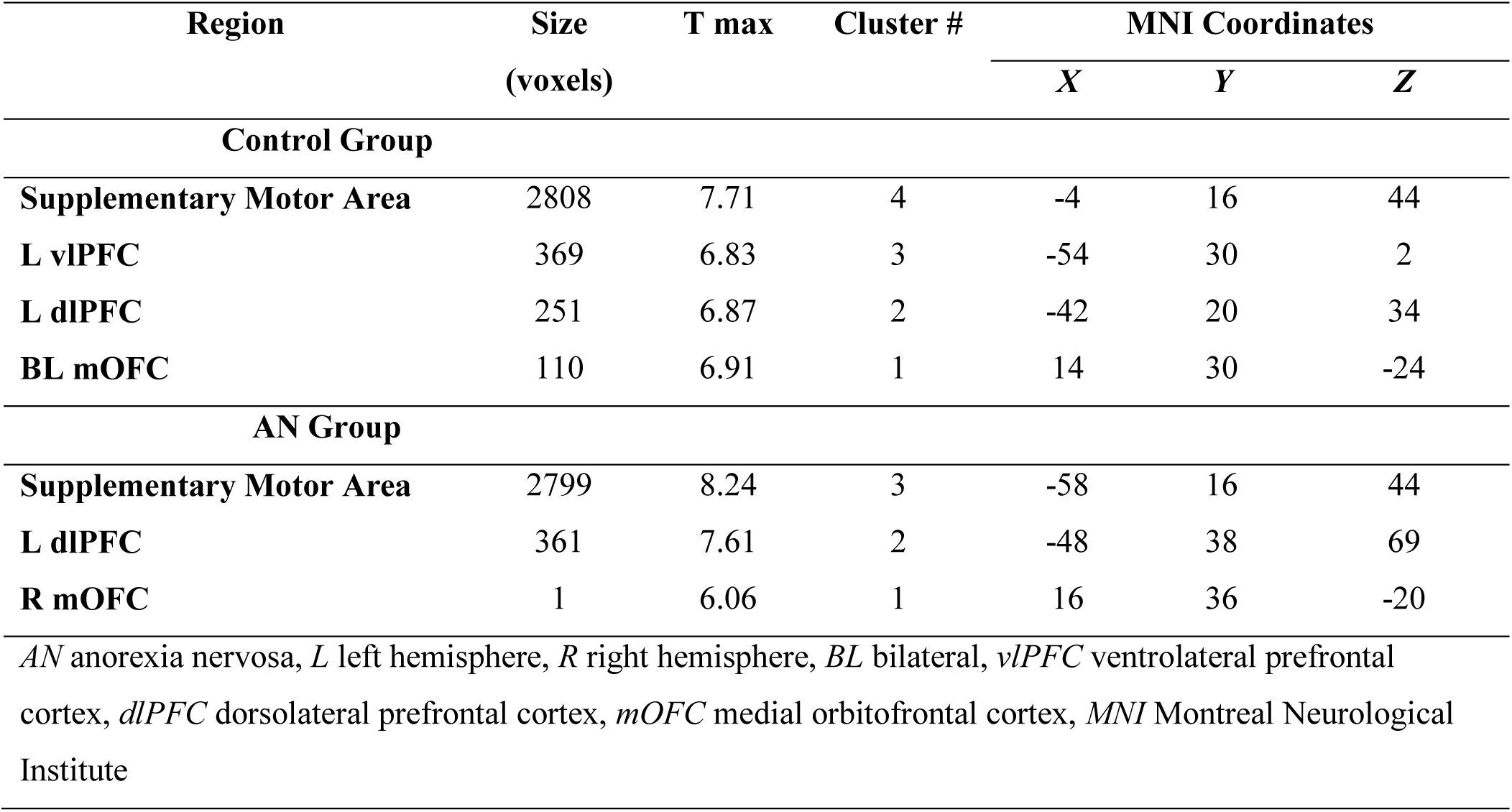
Within-group one-sample *t* – test for anxiety-word rewarded trials for AN and controls.

### 3.3 Associations with clinical variables

The adjusted-BMI LMM found no main effect of time (all *p* > .341), RS (*p* = .183), or their interaction (all *p* > .709). Similarly, with EDE as the dependent variable, no significant effects of time (*p* = .146), RS (*p* = .411), or their interaction (*p* = .214) were detected. Lastly, the LMM with BAS scores found no main effects for time (*p* = .705) or RS (*p* = .310). In summary, the difference in RS between anxiety and neutral reward trials did not predict longitudinal adjusted-BMI or EDE changes, and neutral-word rewarded trial RS were not associated with BAS scores.

Exploratory Pearson and Spearman’s ρ partial correlation analyses examined the relationship between neutral-word rewarded trial RS and clinical measures at baseline, with PDS as a covariate. In the entire sample, a negative relationship between RS and CDRS was found (ρ = −.33, *p* = .028, FDR adjusted *p* = .084) (although not surviving correction for multiple comparisons), suggesting that consistent brain responses may be linked to a lower degree of depressive symptoms. No significant relationship was found between RS and HAM-A (ρ = −.14, *p* = .345, FDR adjusted *p* = .345), or adjusted-BMI (ρ = .24, *p* = .098, FDR adjusted *p* = .147). In the AN group, no significant relationships were detected between RS and EDE (*r* = .05, *p* = .811) or YBC-EDS (*r* = .21, *p* = .326).

## 4 Discussion

This study examined multivariate reward and anxiety brain patterns in adolescents with AN and mildly anxious controls. As hypothesized, individuals with AN had lower RS during neutral-state reward receipt, reflecting less consistent trial-to-trial neural representations within reward-related regions. Within-group searchlight results revealed significant RS clusters in the left mOFC, left NAcc, and left basolateral amygdala in controls but not in AN. Other significant RS clusters appeared in both groups, although with a larger spatial extent in the left vlPFC, left dlPFC, and right mOFC in controls. During reward receipt following an anxiety-inducing word, significant RS clusters were found in the left mOFC and left vLPFC in controls but not AN, although there were no significant group differences, contrary to our predictions.

The difference in RS between anxiety and neutral reward trials did not predict longitudinal adjusted-BMI or eating disorder symptom improvement, and subjective reward responsiveness was not associated with neutral-word rewarded trial RS. Post hoc exploratory analyses identified trend-level relationships between neutral-word reward trials RS and depression, but not anxiety, symptoms.

These results support our hypothesis of lower RS during neutral-word reward trials in the AN group compared with controls. The absence of significant between-group differences for the neutral-word non-rewarded trials suggests that this difference is not just due to general, nonspecific differences in RS during the task. Lower RS may align with univariate literature on attenuated monetary reward responsiveness in AN [30–32]. A prominent theory suggests that this phenomenon is at least partly due to increased top-down cognitive influences related to excessive self-control [33]. Although one study [34] did not find group differences in neural responses during monetary reward receipt in those with acute AN (12-23 years of age), other work [21] found that recovered AN participants (15-28 years of age) had elevated dlPFC activity during monetary reward anticipation, failed to deactivate this region during reward receipt, and displayed greater functional coupling between dlPFC and mOFC. Our multivariate RSA approach provides a different perspective, as it quantifies the consistency of distributed patterns of brain activity across multiple trials, as opposed to averaging brain activity magnitude voxel-by-voxel. Although between-groups searchlight analyses were not significant, the within-group analysis revealed a greater spatial extent of significant RS clusters in controls in the left dlPFC, left vlPFC, and right mOFC, as well as a significant RS cluster that was only present in controls in the left mOFC. Significant trial-by-trial consistency of these frontal areas in controls but not AN was also observed for the neutral-word non-rewarded (right dlPFC), and anxiety-word rewarded trials (left mOFC, left vlPFC). This suggests a more consistent response in controls reflecting more stable processing. In contrast, lower RS in AN may indicate more variable or disrupted neural patterns trial-to-trial.

Further, significant subcortical RS clusters were evident in the left NAcc and left basolateral amygdala in controls but not AN during neutral-state reward receipt. Previous studies have reported different activity patterns in the NAcc of individuals with AN during sucrose reward receipt [35] and with resting-state functional imaging [36]. Additionally, individuals with AN have shown smaller nuclei volume [37, 38] and different connectivity patterns during sucrose reward receipt [39] of the basolateral amygdala. The significant RS clusters in these areas for controls but not in AN suggests the possibility of dynamic pattern differences in reward responses in AN. However, we cannot conclude that significant differences between groups exist for these specific regions, as the between-groups searchlight analyses were not significant.

Following anxiety-induced states, we expected individuals with AN to show a more consistent reward response compared with controls, indicated by higher RS, but this was not observed. This suggests that state anxiety may not differentially affect reward receipt RS between AN and mildly anxious controls. Yet, significant clusters during anxiety-word rewarded trials (left vlPFC and left mOFC), and anxiety-word non-rewarded trials (left basolateral amygdala, right NAcc, and left mOFC) were observed in controls but not in AN. This suggests the possibility that AN may have a pattern of less consistent cognitive control and core reward circuit area reward response, aligning (partially) with findings from the neutral reward condition. However, we may have been underpowered to detect between-group differences for this condition.

Finally, we investigated whether RS response patterns predicted eating disorder-related severity measures longitudinally. RS from anxiety minus neutral reward trials were not associated with changes in adjusted-BMI or EDE, suggesting that that these RS response patterns may not serve as indicators of symptom change. A non-significant main effect of RS also implies that, across time points, those with worse illness phenotypes, as reflected in lower adjusted-BMI and higher EDE scores, do not have more consistent RS. Similarly, neutral-word rewarded trial RS were not associated with BAS scores across baseline and follow-up. Similarly, in our previous work in the same sample (although for a different phase of the task), specific reward and anxiety functional neural connections also did not significantly predict clinical outcomes in AN [40]. Lastly, exploratory correlations showed no significant associations between baseline EDE or YBC-EDS scores and RS.

A somewhat surprising finding was that significant group differences were not observed on the BAS reward responsiveness scale. As discussed in a recent review [9], self-reported reward responsivity in AN is mixed. While a meta-analysis reported lower reward responsivity in acutely ill restricting subtype AN compared to controls and those with other eating disorders [41], other research has found higher trait reward measures (i.e., reward sensitivity, and reward dependence) in AN [42–44]. Compared to norms from the Adolescent Brain Cognition Development (ABCD) general population sample aged 9-20 years (20.22 ± 6.85) [45], both our AN (16.96 ± 2.77) and control group (17.27 ± 2.10) averages were lower. Thus, both groups in our study appear to endorse lower subjective reward responsivity than in the general population. Both groups might share latent contextual factors that account for this. This could include an effect of higher anxiety in both groups than the general population, although if this were the case it would only likely partially account for it since BAS reward responsiveness was similar but the AN group had much higher anxiety ratings, at least as measured on the HAM-A. The lower neural consistency in AN might therefore reflect more subtle or variable processing of rewards that is not captured well by the BAS reward responsiveness self-report measure. Additionally, there were no significant associations between EDE or YBC-EDS scores, suggesting that neural reward response consistency may not be linked to the severity of overall eating disorder symptoms or eating disorder-related obsessive-compulsive symptoms. However, exploratory correlations found trend-level, medium effect size relationships between worse depressive symptoms and lower RS. Notably, the AN group had significantly higher depressive symptoms. The depressive symptom of anhedonia (the loss of pleasure in previously rewarding stimuli) has been found in a meta-analysis to be higher in eating disorder populations than controls [46]. Anhedonia is related to dysregulated reward processing (for a review, see[47]); it is thus possible that anhedonia or other depressive pathophysiological processes in the AN group could partially contribute to, or be a result of, lower neural consistency of reward responses.

Lower reward RS in AN compared to controls may have clinical implications. Adolescents with AN showing less consistent patterns of neural activity during reward receipt may help explain why conventionally rewarding stimuli, like money [30], social engagement [48], and food [49], are perceived as less pleasant or rewarding. Without consistent responses in key reward areas, it is possible that the reliability of experiencing pleasure from these rewarding stimuli may be diminished. Thus, these results may inform newly repurposed Positive Affect Treatment approaches for AN [7], which is a cognitive-behavioral intervention designed to target reward deficits associated with anticipation, experiencing, and learning. As part of monitoring the effects of these interventions, like pleasant event scheduling, investigators could track the consistency of reward responses, perhaps either neurally or with identifying proxy physiological measures that are sensitive to consistency of responses. To establish and validate this approach, this paradigm and analyses may be incorporated into mechanistic clinical trials to understand if the neural reward receipt can be used as a biomarker of response.

Finally, not only were there no significant group differences in reward responses in anxiety-induced states, there were also no significant relationships between reward RS and anxiety symptoms (HAM-A). These results therefore do not provide pre-translational evidence to expect that targeting anxiety symptoms would necessarily improve reward responsiveness.

This study has several limitations. The sample size may have limited our power to detect group differences, particularly for the searchlight analyses. It also limited our ability to conduct sub-analyses of medicated and un-medicated AN groups and to account for comorbidities. While correlation analyses with anxiety and depressive symptoms probed for specific dimensional relationships, they may not fully account for all the effects of comorbid disorders. Replicating this study with a larger sample could help address these issues. Additionally, the study population was at a specific stage of illness recovery— immediately after weight-restoring intensive treatment—so the results might not generalize to other groups.

In conclusion, we found a lower consistency of neural responses for reward receipt in AN compared to mildly anxious controls. Potentially accounting for this is low consistency of responses in core nodes of the reward circuit as well as in reward-related prefrontal cognitive control regions in AN. However, induced anxiety-states did not affect reward responses differentially in AN and controls. The results from these analyses have implications for future research and for the ongoing development of treatment approaches targeted towards improving reward processing in AN.

## Acknowledgments

The authors declare no conflicts of interest. This study was supported by the National Institute of Mental Health (R01MH105662 to JDF)

## Author Contributions

JDF and MS concepted and designed the study. HJP and JDF wrote the first draft of the manuscript. NR, HJP, and JDF contributed to the statistics and analyses of the paper. All authors worked on the final draft and gave their approval for submission.

## Data Availability

The data that support the findings of this study are openly available in the NIMH Data Archive at https://nda.nih.gov/, collection ID 2565.

## Notes

### Competing Interest Statement

The authors have declared no competing interest.

