## Supplementary Materials for "Multivariate Neural Patterns of Reward and Anxiety in Adolescents with Anorexia Nervosa"

### Title

**Contents:**

**Supplementary Methods**


1. Inclusion / Exclusion criteria
2. Clinical assessments
3. MRI data acquisition
4. Functional data preprocessing
5. Calculation of beta values, additional preprocessing, and missing data
6. FMRI paradigm and word selection
7. Region of interest creation
8. Details on exploratory anxiety-word post-reward analysis


**Supplementary Tables**


**Table S1** Within-group one-sample t-test results for anxiety-word non-rewarded trials
**Table S2** Within-group one-sample t-test results for neutral-word non-rewarded trials
**Table S3** Within-group one-sample t-test results for anxiety-word post-rewarded trials


**Supplementary Figures**


**Figure S1** Within-group one-sample t-test results from searchlight analysis within the reward mask for the anxiety-word and neutral-word non-rewarded trials
**Figure S2** Within-group one-sample t-test results from searchlight analysis within the anxiety mask for the anxiety-word post-rewarded trials


**Supplementary References**

### 1. Inclusion / Exclusion criteria

Participants with AN met the DSM-5 criteria for the restricting type of AN. Inclusion criteria for the AN group were: Ages 10–19; met DSM-5 criteria for Anorexia Nervosa, Restricting Type, within the previous 6 months; completed usual treatment in an inpatient, residential, or partial hospitalization program (2–5 times/week) involving psychotherapy and dietary monitoring within the previous 3 weeks; were either un-medicated or taking a stable dose of serotonin reuptake inhibitors for at least 8 weeks; and finally, they could not be on any other psychotropic medications except for a short half-life sedative/hypnotic for insomnia or a short half-life benzodiazepine for anxiety, not exceeding three doses per week and not taken on scan days. Exclusion criteria for the AN group included a lifetime history of bipolar disorder, psychotic disorders, attention-deficit hyperactivity disorder, or current post-traumatic stress disorder.

Inclusion criteria for control participants were females aged 10–19 who scored at least 0.5 standard deviations higher than population norms on the anxiety portion of the Depression Anxiety Stress Scale (DASS-21). This helped us better examine the relationship between anxiety and brain function and structure across AN and control groups. Exclusion criteria for control participants were any DSM-5 diagnosis, assessed with the MINI KID 7.0.2, and use of any psychiatric medication.

Exclusion criteria for all participants were: current substance abuse or dependence, including nicotine; pathological gambling, as assessed with the South Oaks Gambling Screen; current medical and neurological disorders (e.g., diabetes, hypertension, seizure disorders, migraine headaches) requiring treatment at the time of the experiment; pregnancy; major medical disorders that could affect cerebral metabolism, such as diabetes or thyroid disorders; current risk of suicide with a plan and intent; a Children's Depression Rating Scale Revised (CDRS-R) score >75 (extremely ill) or major depressive disorder with psychotic features; ferromagnetic metal implants or devices (e.g., electronic implants, infusion pumps, aneurysm clips, metal fragments, metal prostheses, joints, rods, or plates); BMI ≥ 25 (overweight); and visual acuity worse than 20/35 for each eye, as determined by a Snellen close vision chart (with corrective lenses allowed).

### 2. Clinical assessments

Clinical evaluations for all participants were conducted by licensed psychiatrists or psychologists experienced with this population. Primary or comorbid diagnoses were screened using the Mini-International Neuropsychiatric Interview (MINI KID 7.0.2) [1]. To measure depression severity, the Children's Depression Rating Scale™, Revised (CDRS™-R) [2] and the depression subscale of the Depression Anxiety Stress Scale (DASS-21) [3] were administered. We also administered the Behavioral Inhibition System/Behavioral Activation System (BIS/BAS) [4] questionnaire (for this analysis we focused on the BAS Reward Responsiveness scale, as the construct it measures most closely corresponds to the responsivity to reward portion of the fMRI task). The Hamilton Anxiety Rating Scale (HAMA) [5] measured past week anxiety. Additionally, participants rated their level of anxiety immediately after scanning on a Likert scale from 0 to 10. To evaluate the severity of AN symptoms, the Eating Disorder Examination (EDE) and the Yale-Brown-Cornell Eating Disorder Scale (YBC-EDS) [6] were administered.

### 3. MRI acquisition

MRI data were acquired on a 3T Siemens PRISMA scanner using a 64-channel coil. Echoplanar images (EPI) were collected with the following parameters: repetition time (TR) of 1000 ms, echo time (TE) of 33 ms, flip angle of 80°, isotropic voxel size of 2 $mm^3$, multiband acceleration factor of 5, field of view of 208 mm, 487 volumes, and 60 slices. Structural MRI for registration purposes utilized a T1-weighted MPRAGE sequence with a TR of 2300 ms, TE of 2.99 ms, and isotropic voxel dimensions of 0.8 $mm^3$.

### 4. Functional data preprocessing

FSL (FMRIB's Software Library, www.fmrib.ox.ac.uk/fsl) with FEAT version 6.0 (FMRI Expert Analysis Tool) was used to preprocess the anxiety and neutral fMRI runs of each participant. This included motion correction (MCFLIRT) and temporal filtering. The unsmoothed, native functional space fMRI runs from each participant were then forwarded to the RSA preprocessing pipeline (see below). Additionally, T1-weighted scans were brain-extracted using BET and used for standard space registration with FLIRT before conducting RSA.

### 5. Calculation of beta maps and missing data

We used the same protocol for calculating beta maps as in our previous RSA study [7]. Briefly, we extracted the relevant time series corresponding to the reward and non-reward period of the anxiety and neutral runs, as well as the anxiety word period after rewarded trials, using custom MATLAB scripts. The preprocessed time series data in native space was then forwarded to AFNI [8] and deconvolved using the 3dDeconvolve command. We conducted least-square-sum estimates of beta values using 3dLSS for each reward/word period within the respective run. The approximately 30 single-trial beta maps from each participant for the reward/word periods were transformed into MNI standard space using the previously calculated transformation matrices within FSL and then forwarded to the RSA pipeline. Time points with movement outliers, identified using FSL's motion outlier tool (fsl_motion_outliers) with the default threshold, and were excluded from further analysis based on the DVARS metric [9].

Three participants had missing values for the PDS data. To address this, we imputed the missing data by using the mean value of other participants in the same age range (±1 year) within the respective group.

### 6. FMRI paradigm

Before the scans, participants were told that they were going to play a game in which they can earn $10 for correct responses. They were required to classify 6 unique colored fractal images as belonging (arbitrarily) to ‘Group 1’ or ‘Group 2’. At the beginning of each trial, 1 of 6 randomly generated fractals were presented for 2000 ms. Participants were instructed to push the right or left button to guess if it belongs (arbitrarily) in “Group 1” or “Group 2.” After button pressing, there was an inter-stimulus-interval jittered randomly between 2500 and 1250 ms, followed by a word stimulus (anxiety-evoking or neutral) presented for 2000 ms. Then, after another 250 ms blank screen inter-stimulus-interval, participants were given feedback about receiving a reward or not receiving a reward. The chance of receiving a reward in each block was random (50% probability), and therefore, did not invoke learning. Another inter-stimulus-interval randomly jittered between 1250 and 2500 ms appeared prior to the next fractal. There were 60 randomized trials presented in total, with the 20 anxiety-evoking words presented 3 times each. Each trial lasted 6.43 ms on average, and the total run length was 498 seconds.

Before scanning, all participants viewed 150 words on a computer screen and asked “How anxious does this word make you feel?”. Words were rated on a scale from 1-9 (with 9 being the highest). The words were sourced from three lists: 50 anorexia-related anxiety words (e.g., fat, diet), 50 general anxiety words (e.g., harm, pain), and 50 neutral words (e.g., bus, shop), all adapted from previous studies [10-14]. The words were presented in a random order. For the AN group, only anorexia-related anxiety words were used, while for the control group, only general anxiety words were presented. Each participant's top 20 words with the highest anxiety ratings were selected for the anxiety run. The neutral words for both groups were chosen from the neutral list, specifically the 20 words rated as least anxiety-provoking. Additionally, we ensured that the overall number of syllables did not significantly differ between the two lists for each participant. If there were significant differences, the next-highest (anxiety) or next-lowest (neutral) word was selected until the syllable count was balanced. Participants were randomly assigned to either the anxiety or neutral word task at the first scan, with the other task occurring during the second scan.

### 7. Region of interest creation

Probabilistic atlases thresholded at 50% were used to extract the basolateral amygdala (Harvard-Oxford), mOFC (aal2) and SMA (aal2). To create masks corresponding to reward-related cognitive control for the vlPFC and dlPFC, the left vlPFC mask was created based on the Harvard-Oxford atlas for the pars triangularis, again thresholded at 50%. The dlPFC was created by first merging the superior- and middle-frontal probabilistic maps from the Harvard-Oxford atlas, then, the voxels overlapping these merged maps and the association map generated from the Neurosynth fMRI meta-analytic database (neurosynth.org) using the search term "DLPFC" were used as the final mask. The Pauli and colleagues [28] atlas was used for the VTA and NAcc, but given these are small, they were thresholded at values of 0.001. All regions were then merged into a single ROI mask.

### 8. Details on exploratory anxiety-word post-reward trials analysis

As research [15] has shown how a positive state (e.g., positive affect after acute exercise) is associated with reductions in state anxiety, we conducted an exploratory analysis, testing if a subsequent anxiety state is influenced by a preceding reward receipt, as we reasoned that being in a rewarded state could engage overlapping anxiety and reward circuitry, thereby diminishing anxiety responsiveness. Thus, our exploratory hypothesis was that AN participant's response to anxiety provocation in anxiety regions for trials after reward receipt will be more consistent than controls.

The anxiety ROI mask used for this analysis comprised the same areas as in our previous studies [7, 16], with the exception of the centromedial amygdala nuclei instead of the entire amygdala. The other areas included the anterior cingulate cortex, insula, medial prefrontal cortex, ventral tegmental area, and bed nucleus of the stria terminalis.

ANCOVA found no significant group differences in representational similarity in the anxiety ROI mask for anxiety-word trials post-reward ($F_{(1, 44)} = 0.05$, $p = .822$). Between-group searchlight analyses did not reveal any significant differences between groups, and MANOVA did not find that any one region contributed more strongly to results. Within-group searchlights found significant clusters of representational similarity for the AN group which had much larger spatial extents than controls in the right centromedial amygdala, and anterior portions of the left insula (Figure S2, Table S3).

This exploratory analysis examined whether anxiety-word post-reward trial RS was higher in AN than controls. Although no group differences emerged, within-group searchlight analyses revealed significant RS in the right centromedial amygdala and left anterior insula in AN, with a larger spatial extent and magnitude compared to controls. The anterior insula, involved in anxiety and fear regulation[17], may play a crucial role in eating disorders and is associated with structural and functional differences compared to controls [18-20]. The centromedial amygdala is highly sensitive to negative emotional stimuli [21], and amygdala hyperreactivity to a range of stimuli [22], along with altered nuclei volume in AN [23, 24] have been reported, although these studies focused on the amygdala broadly rather than the centromedial nuclei specifically. These findings suggest consistent engagement of the anterior insula and centromedial amygdala in AN during anxiety word processing. However, as with our analysis testing if anxiety-induced states influence reward receipt RS, conclusions about the influence of prior reward receipt on these patterns are difficult to infer. From our results, there is some preliminary evidence that anxiety-word processing may differ in terms of which regions contribute more strongly to representational similarity in AN, but determining if this is due to prior reward receipt or just differences in processing anxiety stimuli may benefit from a transfer learning approach in future research.

**Table S1** Within-group one-sample t-test results for anxiety-word non-rewarded trials

| Region | Size (voxels) | T max | Cluster # | MNI Coordinates | | |
|---|---|---|---|---|---|---|
| | | | | ***X*** | ***Y*** | ***Z*** |
| **Control Group** | | | | | | |
| **Supplementary Motor Area** | 2471 | 9.44 | 7 | -4 | 8 | 46 |
| **L vlPFC/dlPFC** | 646 | 7.09 | 6 | -52 | 30 | 6 |
| **R mOFC** | 83 | 8.20 | 5 | 20 | 28 | -22 |
| **L Basolateral Amygdala** | 46 | 7.42 | 4 | -24 | -8 | -26 |
| **L mOFC** | 6 | 7.42 | 3 | -14 | 20 | -20 |
| **R NAcc** | 2 | 7.83 | 2 | 12 | 6 | -10 |
| **AN Group** | | | | | | |
| **Supplementary Motor Area** | 2646 | 10.20 | 7 | 0 | 18 | 44 |
| **L vlPFC/dlPFC** | 249 | 7.21 | 6 | -50 | 22 | 4 |
| **L dlPFC** | 78 | 5.93 | 5 | -38 | 34 | 34 |
| **R Basolateral Amygdala** | 19 | 7.43 | 4 | 20 | -2 | -26 |
| **R mOFC** | 18 | 7.15 | 3 | 18 | 30 | -24 |
| **L dlPFC** | 3 | 5.38 | 2 | -46 | 12 | 10 |

Note. L = left hemisphere, R = right hemisphere, vlPFC = ventrolateral prefrontal cortex, dlPFC = dorsolateral prefrontal cortex, mOFC = medial orbitofrontal cortex, NAcc = nucleus accumbens, MNI = Montreal Neurological Institute, AN = anorexia nervosa

**Table S2** Within-group one-sample t-test results for neutral-word non-rewarded trials

| Region | Size (voxels) | T max | Cluster # | MNI Coordinates | | |
|---|---|---|---|---|---|---|
| | | | | ***X*** | ***Y*** | ***Z*** |
| **Control Group** | | | | | | |
| **Supplementary Motor Area** | 2597 | 8.89 | 7 | 0 | 16 | 44 |
| **L vlPFC** | 376 | 6.39 | 6 | -50 | 32 | 6 |
| **L dlPFC** | 172 | 6.15 | 5 | -42 | 26 | 32 |
| **R mOFC** | 152 | 6.86 | 4 | 14 | 36 | -24 |
| **L mOFC** | 112 | 6.81 | 3 | -14 | 36 | 22 |
| **L Basolateral Amygdala** | 44 | 7.03 | 2 | -24 | -8 | -24 |
| **R dlPFC** | 21 | 7.13 | 1 | 38 | 34 | 36 |
| **AN Group** | | | | | | |
| **Supplementary Motor Area** | 2180 | 8.94 | 6 | -4 | 6 | 46 |
| **L vlPFC** | 198 | 6.91 | 5 | -52 | 24 | 2 |
| **L mOFC** | 145 | 7.82 | 4 | -16 | 16 | -18 |
| **L dlPFC** | 132 | 7.41 | 3 | -46 | 24 | 32 |
| **R mOFC** | 92 | 6.09 | 2 | 22 | 32 | -18 |
| **R Basolateral Amygdala** | 1 | 6.09 | 1 | 28 | -6 | 22 |

Note. L = left hemisphere, R = right hemisphere, vlPFC = ventrolateral prefrontal cortex, dlPFC = dorsolateral prefrontal cortex, mOFC = medial orbitofrontal cortex, NAcc = nucleus accumbens, MNI = Montreal Neurological Institute, AN = anorexia nervosa

**Table S3 Within-group one-sample *t* – test for anxiety-word post-rewarded trials for AN and controls**

| Region | Size (voxels) | T max | Cluster # | MNI Coordinates | | |
|---|---|---|---|---|---|---|
| | | | | ***X*** | ***Y*** | ***Z*** |
| **Control Group** | | | | | | |
| **R ACC** | 1364 | 7.27 | 7 | 4 | 36 | -6 |
| **R Insula** | 196 | 7.17 | 6 | 40 | -4 | 6 |
| **L PFC** | 122 | 5.42 | 5 | -4 | 44 | -20 |
| **L Insula** | 84 | 6.19 | 4 | -40 | -6 | -10 |
| **R Insula** | 36 | 5.87 | 3 | 36 | 12 | -14 |
| **R Centromedial Amygdala** | 2 | 5.93 | 2 | 28 | -4 | -12 |
| **R Centromedial Amygdala** | 2 | 5.80 | 1 | 28 | -6 | -8 |
| **AN Group** | | | | | | |
| **R ACC** | 1387 | 8.24 | 7 | -6 | 38 | -2 |
| **L Insula** | 385 | 8.42 | 6 | -38 | -2 | -12 |
| **R Insula** | 360 | 7.51 | 5 | 42 | 4 | -10 |
| **L PFC** | 149 | 5.47 | 4 | -4 | 38 | -18 |
| **R Centromedial Amygdala** | 23 | 6.76 | 3 | 22 | -10 | -6 |
| **L PFC** | 11 | 5.68 | 2 | -2 | 54 | -18 |
| **R Centromedial Amygdala** | 3 | 6.00 | 1 | 28 | -12 | -6 |

*AN* anorexia nervosa, *L* left hemisphere, *R* right hemisphere, *ACC* anterior cingulate cortex, *PFC* prefrontal cortex, *MNI* Montreal Neurological Institute

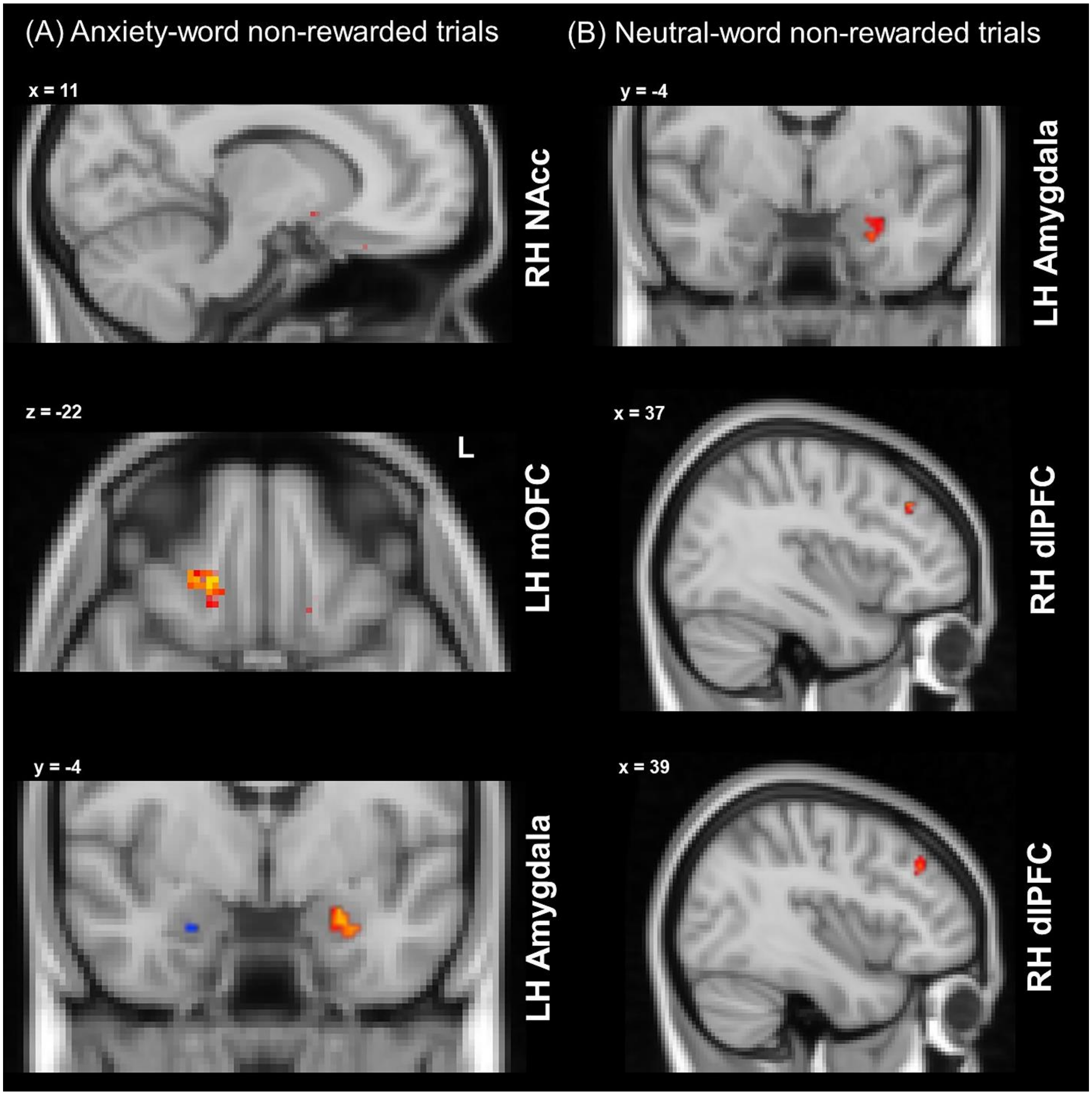


*Figure S1.* Within-group one-sample t-tests for the (A) anxiety-word non-rewarded trials representational similarity from the searchlight analysis within the reward mask, and (B), neutral-word non-rewarded trials after reward receipt representational similarity from the searchlight analysis within the anxiety mask. Within-group results are overlaid: red/yellow clusters depict significant areas in controls, while blue clusters are significant in AN ($p$ < .05, corrected).

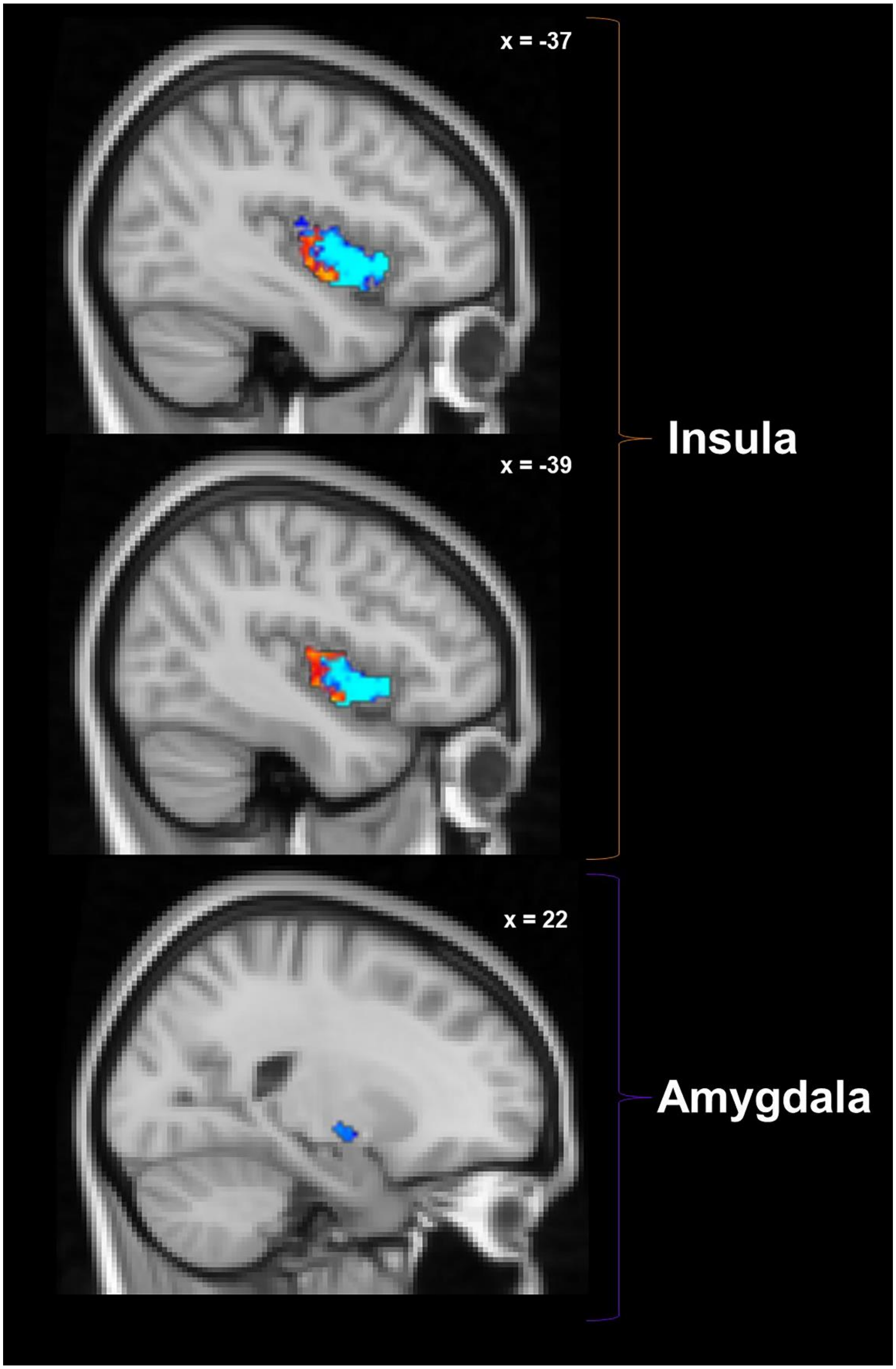


*Figure S2*. Within-group one-sample t-tests for the anxiety-word after rewarded trials representational similarity from the searchlight analysis within the anxiety mask. Within-group results are overlaid: red/yellow clusters depict significant areas in controls, while blue clusters are significant in AN ($p < .05$, corrected).
